# STEQ: A statistically consistent quartet distance based species tree estimation method

**DOI:** 10.64898/2026.02.27.708511

**Authors:** Prottoy Saha, Anik Saha, Mrinmoy Saha Roddur, Sagor Sikdar, Nayeem Hasan Anik, Rezwana Reaz, Md. Shamsuzzoha Bayzid

## Abstract

Accurate estimation of large-scale species trees from multilocus data in the presence of gene tree discordance remains a major challenge in phylogenomics. Although maximum likelihood, Bayesian, and statistically consistent summary methods can infer species trees with high accuracy, most of these methods are slow and not scalable to large number of taxa and genes. One of the promising ways for enabling large-scale phylogeny estimation is distance based estimation methods. Here, we present STEQ, a new statistically consistent, fast, and accurate distance based method to estimate species trees from a collection of gene trees. We used a quartet based distance metric which is statistically consistent under the multi-species coalescent (MSC) model. The running time of STEQ scales as 𝒪 (*kn*^2^ log *n*), for *n* taxa and *k* genes, which is asymptotically faster than the leading summary based methods such as ASTRAL. We evaluated the performance of STEQ in comparison with ASTRAL and wQFM-TREE – two of the most popular and accurate coalescent-based methods. Experimental findings on a collection of simulated and empirical datasets suggest that STEQ enables significantly faster inference of species trees while maintaining competitive accuracy with the best current methods. STEQ is publicly available at https://github.com/prottoysaha99/STEQ.

## 1 Introduction

Estimation of the species tree from multiple genes sampled throughout the whole genome has become a fundamental task in comparative and evolutionary biology. However, accurate species tree estimation from multiple genes is not easy since true gene trees can differ from each other, a phenomenon known as gene tree heterogeneity or gene tree discordance [19]. Incomplete lineage sorting (ILS), modeled by multi-species coalescent (MSC) [10,11], is one of the dominant reasons for gene tree heterogeneity.

The standard approach to estimate species trees from multi-locus data is called concatenation or combined analysis. This approach concatenates gene sequence alignments into a single supergene alignment, and then estimates species tree from this supergene alignment [8]. However, concatenation can be statistically inconsistent [31], and positively misleading [13,6,15,5]. To address the discordance among gene trees, “summary methods” have been developed which compute gene trees from different loci and then use the set of gene trees to infer a species tree. Many of the summary methods are statistically consistent under the MSC model, including ASTRAL [22,23,37], MP-EST [17], NJst [16], BUCKy [14], GLASS [24], STEM [12], SNAPP [3], STELAR [9], SVDquartets [4], STEAC [18] and ASTRID [35]. ASTRAL, currently the most widely used quartet-based method, infers species trees by maximizing the number of gene tree quartets that are consistent with the species tree. An alternative strategy (e.g., Quartet Max-Cut (QMC) and Quartet Fiduccia–Mattheyses (QFM) families of methods [33,27,2,20]) first infers quartets (often with weights) for all four-taxon subsets and then amalgamates them into a single, coherent species tree. These methods are highly accurate, but they face scalability limitations due to its high computational cost of explicitly generating and weighting quartets. But faster versions have recently been developed (TREE-QMC [7] and wQFM-TREE [25]) that apply the QMC and QFM heuristic directly to gene trees.

Previous works on quartet-based distance include “anchor” based computation of pairwise distances [32], which chooses two taxa as anchors to compute the probability of observing a quartet. Yourdkhani et al. [36] estimated species trees from weighted quartets or rooted triples derived from gene trees. They defined inter-taxon distances based on the internal edge weights of these quartets/triplets. Rhodes proposed a method [28], which determines the set 𝒬 of *dominant* quartets (i.e., most frequent quartets) for each subset of four taxa; and then for each pair (*x, y*) of taxa, it counts the number of quartets in 𝒬 separating *x* and *y* to build distance matrices. MSCquartets [29], an R package, estimates species tree using a distance-based method where inter-taxon distance matrices are constructed from the quartet count concordance factor. However, [36,29,28] face inherent scalability limitations due to their high computational complexity, particularly with the *O*(*n*^4^*k*) quartet enumeration step, where *n* is the number of taxa and *k* is the number of gene trees. Other notable distance-based methods include ASTRID [35], which is a simple modification to NJst [16].

In this paper, we present STEQ (**S**pecies **T**ree **E**stimation using **Q**uartet distance), a new statistically consistent, ILS-aware, and scalable method for species tree estimation. STEQ is a distance based method, where distance between two species is calculated as the average number of induced quartets in the gene trees where the two species lie on the different sides of the bi-partition defined by the internal edge of the quartet. STEQ computes the distance matrix directly from gene trees without explicitly enumerating the quartets. We also introduced a novel normalization technique for improved accuracy. After computing the dis-tance matrix, STEQ uses FASTME or BioNJ (if there are missing taxa in the gene trees) to estimate the species tree. With a time complexity of *O*(*n*^2^*k* log *n*) (where *n* is the number of taxa and *k* is the number of gene trees), STEQ is both asymptotically and empirically faster than wQFM-TREE and ASTRAL, particularly on larger datasets. Experimental results demonstrate that STEQ achieves comparable or superior accuracy to wQFM-TREE and ASTRAL across a wide range of datasets, highlighting its suitability for accurately analyzing thousands of taxa and gene trees.

## 2. Methods

### 2.1 Quartet distance between two species

STEQ takes as input a set 𝒢= {*gt*_1_, *gt*_2_, …, *gt*_*k*_} of *k* gene trees on a set 𝒳= {*x*_1_, *x*_2_, …, *x*_*n*_} of *n* taxa. STEQ can handle both rooted and unrooted gene trees and it allows incomplete gene trees meaning that they can have missing taxa. Let *L*(*t*) denote the set of taxa present in a tree *t*. Then *L*(*gt*_*i*_) ⊆ 𝒳, and we call a gene tree incomplete when *L*(*gt*_*i*_) ⊊ 𝒳. A *quartet q* is an unrooted tree with four taxa. We denote by *q* = *ab* |*cd*, where *a* and *b* are separated from *c* and *d* by an edge. For a quartet *q* = *ab*|*cd*, we call *a* and *b* (also *c* and _(_*d*) _t_are on the *same side* of *q*. Note that a tree *gt* with |*L*(*gt*_*i*_)| = *n*_*i*_ taxa induces 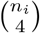 quartets.

STEQ defines the quartet distance 𝒬 𝒟_*gt*_(*x, y*) between two taxa *x* and *y* in a gene tree *gt* as the number of quartets in *gt* that contain both *x* and *y*, where *x* and *y* lie on different sides. If one or both of *x* and *y* are not present in *t*, 𝒬 𝒟_*gt*_(*x, y*) is set to zero. Distance 𝒬 𝒟 (*x, y*) between species *x* and *y* is defined as the average quartet distance between the gene copies sampled from species *x* and species *y* across all gene trees. To calculate 𝒬 𝒟 (*x, y*), we consider only those gene trees that contain both *x* and *y*. Let *N*_*gt*_(*x, y*) denote the number of gene trees that contain both *x* and *y*. Then 𝒬 𝒟 (*x, y*) is defined as follows.

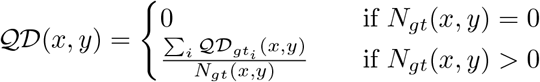

We define an *n* × *n* distance matrix ℳ, where ℳ (*i, j*) is defined as,

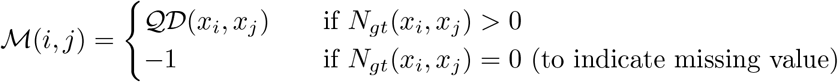

Finally, we estimate the species tree on the distance matrix ℳ using variants of neighbor joining (BioNJ or FastME).

### 2.2 Computing the quartet distance

Fundamental to our method is the ability to efficiently calculate the quartet distance between two taxa in a gene tree. We now define some preliminaries required to define the quartet distance. Let *gt* be a gene tree and *u* be an arbitrary internal node. Note that three edges are incident on any internal node and thus *u* defines a tripartition *A* |*B*| *C*. For any quartet *q* = *ab* |*cd* in *gt, q* maps to two internal nodes *u* and *u*^′^. *u* is the node where the path from *a* to *c* (or *d*) intersects with the path from *b* to *c* (or *d*) for the first time. Similarly, the paths from *c* to *a* (or *b*) and *d* to *a* (or *b*) first intersect with each other at *u*^′^ (see Supplementary Fig. S1). This concept of mapping was previously described in [22]. It is easy to see that the number of quartets mapped to an internal node *u* with a tripartition *A*|*B*|*C* is,

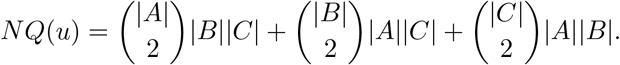

Since each quartet is mapped to two internal nodes, total number of induced quartets in a tree is given by 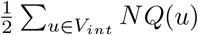, where *V*_*int*_ is the set of internal nodes in the tree.

We now describe how to calculate the quartet distance 𝒬 𝒟_*gt*_(*x, y*) between two taxa *x* and *y* in *gt*. Let *U*_*xy*_ be the set of internal nodes on the path from *x* to *y*, excluding *x* and *y*. It is easy to see that for any internal node *u* ∉ *U*_*xy*_, the quartets mapped to *u* will have *x* and *y* on the same side (see Fig. 1a), and thus will not contribute towards 𝒬 𝒟_*gt*_(*x, y*). Therefore, we need to consider only the internal nodes in *U*_*xy*_.

**Fig. 1.**
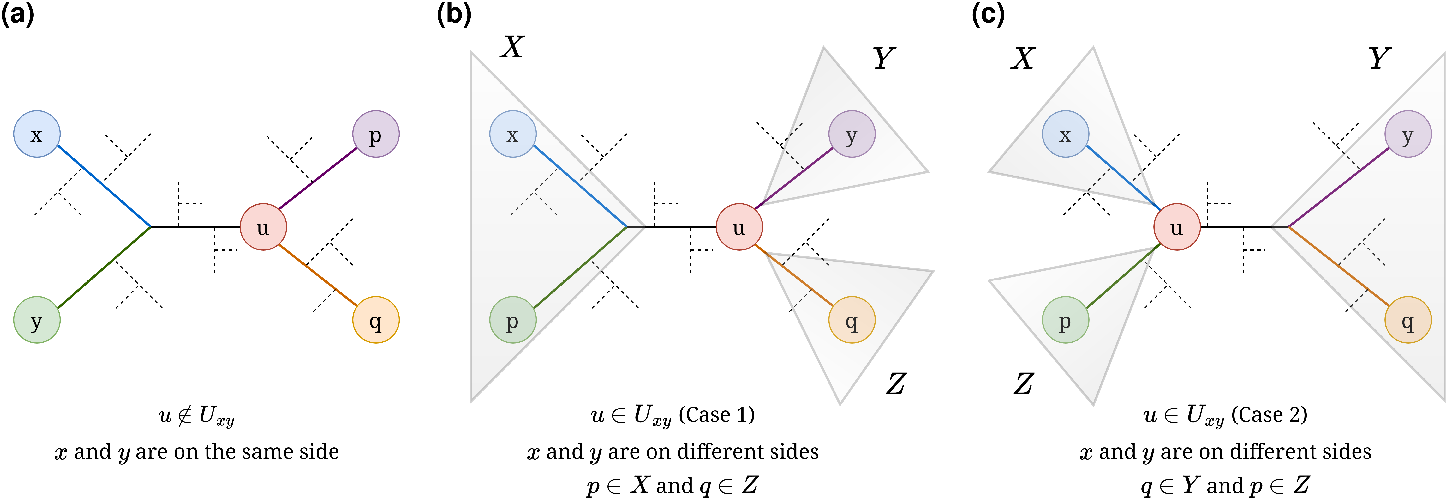
Placements of *u* with respect to *x* and *y*. (a) *x* and *y* lie on the same side, *u* ∈*/ U*_*xy*_ (b)-(c) *x* and *y* lie on different sides, *u* ∈ *U*_*xy*_ (b) Case 1 where *p* ∈ *X* and *q* ∈ *Z* (c) Case 2 where *q* ∈ *Y* and *p* ∈ *Z*

Let 𝒬 𝒟_*u*_(*x, y*) denote the number of quartets that are mapped to *u* ∈ *U*_*xy*_ and have *x* and *y* on two different sides. We denote by *X* |*Y* |*Z* the tripartition defined by *u* where *x* ∈ *X* and *y* ∈ *Y*. Let *x, y, p, q* be the four taxa in a quartet that has been mapped to *u*. We want to analyze when *x* and *y* will be on the opposite side of the quartet. Without loss of generality, let *p* be closer to *x* and *q* be closer to *y* yielding the quartet topology (*px*| *qy*). Then either of the following two conditions holds (see Fig. 1b-c).

1. If *u* is the first common node of the paths from *y* and *q*: *p* ∈ *X* and *q* ∈ *Z*
2. If *u* is the first common node of the paths from *x* and *p*: *q* ∈ *Y* and *p* ∈ *Z*

Thus, 𝒬 𝒟_*u*_(*x, y*) can be calculated as follows.

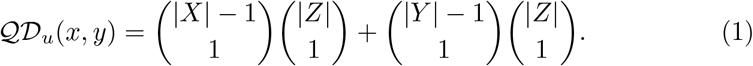

Therefore, 𝒬 𝒟_*gt*_(*x, y*) can be calculated as follows. Note that the division by 2 is due to the fact that each quartet is mapped to two different internal nodes.

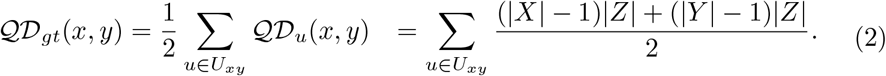

### 2.3 Statistical consistency

We now prove that STEQ is statistically consistent under the MSC model.

#### Theorem 1.

*The quartet distance is additive on a gene tree*.

*Proof*. To prove that quartet distance is additive, it is sufficient to prove that the four-point condition holds for any arbitrary set of four species. Let *gt* be a gene tree for a particular gene sampled from *n* species. Consider four arbitrary taxa *a, b, c, d* in *gt*. Without loss of generality, assume that *a* and *b* are more closely related to each other than they are to *c* and *d* in *gt*. That means *q* = *ab* |*cd* is the quartet induced by these four taxa in *gt*. Let *q* map to two internal nodes *u* and *u*^′^ (see Supplementary Fig. S1).

To verify the four-point condition, we need to compare sums of pairwise quartet distances among *a, b, c, d*. The key idea is that 𝒬 𝒟_*gt*_(*x, y*) is a sum of contributions from internal nodes on the path *U*_*xy*_. For the quartet *ab* |*cd*, all cross-pair paths share the same central segment between the two mapping nodes *u* and *u*^′^; the remaining segments are leaf-specific. Thus, we partition *U*_*ac*_ into subpaths to reuse common summands and thereby simplify the four-point comparison.

We now compute the pairwise distances between *a, b, c* and *d*. The distance 𝒬 𝒟_*gt*_(*a, c*) between *a* and *c* can be calculated as follows according to Equation 2.

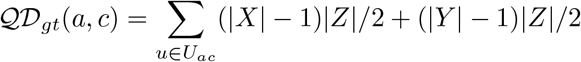

*U*_*ac*_ contains all the internal nodes on the path from *a* to *c* in *gt*. We divide *U*_*ac*_ into 5 subsets: i) 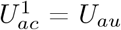 (internal nodes from *a* to *u* excluding *u*), ii) 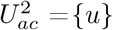, iii) 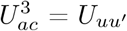 (internal nodes from *u* to *u*^′^ excluding *u* and *u*^′^), iv) 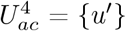, and v) 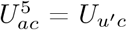 (internal nodes from *u*^′^ to *c* excluding *u*^′^)

Thus, 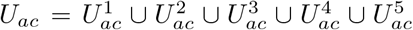. Let the number of taxa descending from the internal nodes on the path from *a* to *u* (excluding *u*) is *A*, and *B* be the number of taxa descending from the internal nodes on the path from *b* to *u* (excluding *u*). Similarly, we can define *C* and *D* with respect to the other mapping node *u*^′^. We now define the following distances for computational convenience

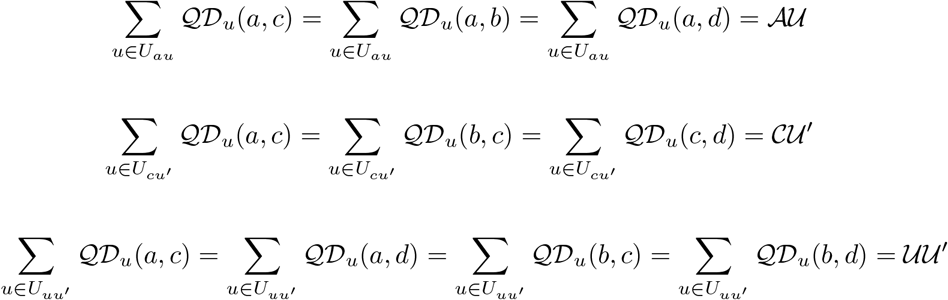

Similarly, we can define ℬ 𝒰 and 𝒟 𝒰^′^. Now,

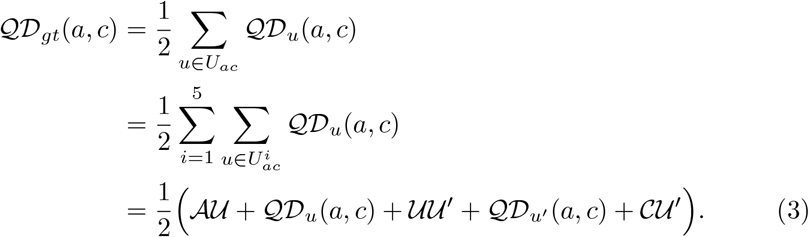

Here, according to Equation 1,

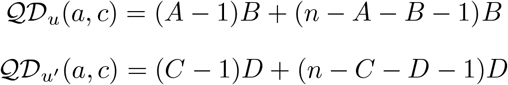

Therefore, substituting into Equation 3,

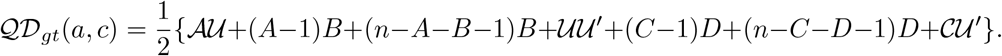

Similarly,

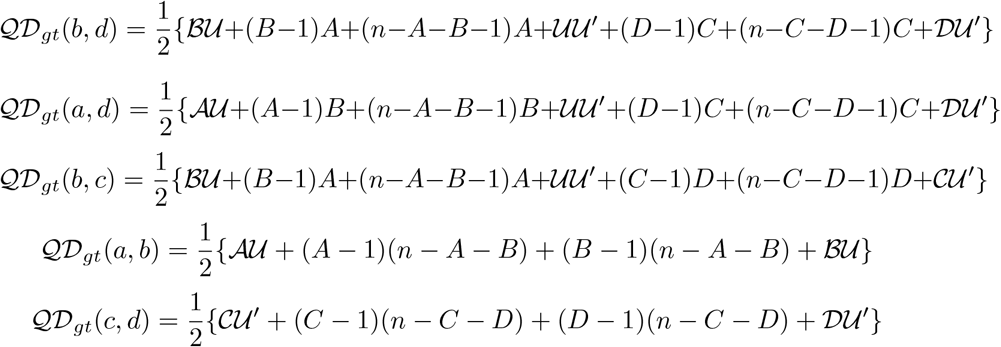

Now, we compute the value of 𝒬 𝒟_*gt*_(*a, c*) + 𝒬 𝒟_*gt*_(*b, d*).

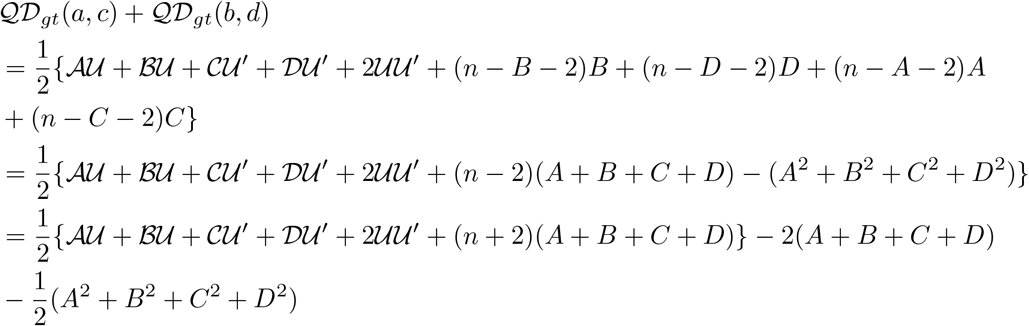

Similarly, it is easy to see that,

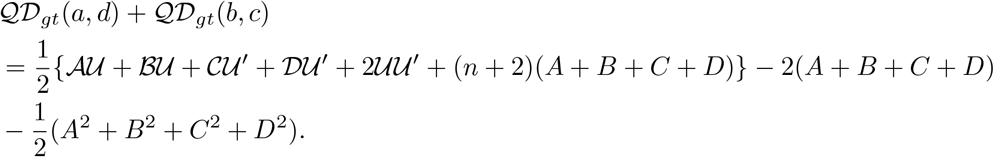

Therefore,

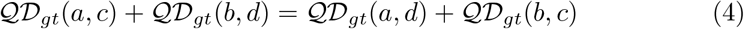

We now compute the value of 𝒬 𝒟_*gt*_(*a, b*) + 𝒬 𝒟_*gt*_(*c, d*).

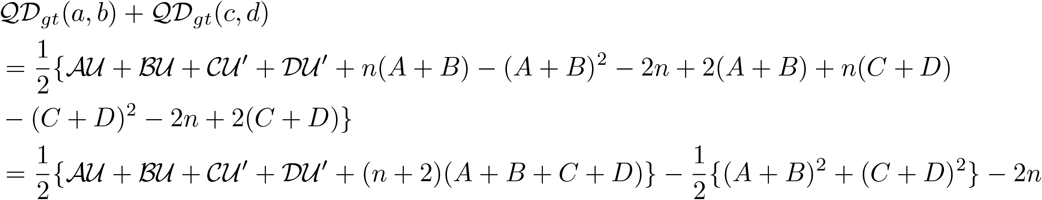

Note that 𝒰 𝒰 ≥ ′ 0 and *A* + *B* + *C* + *D ≤ n*. Moreover, since *A >* 0, *B >* 0, *C >* 0 and *D >* 0, (*A* + *B*)^2^ + (*C* + *D*)^2^ *>* (*A*^2^ + *B*^2^ + *C*^2^ + *D*^2^). Therefore,

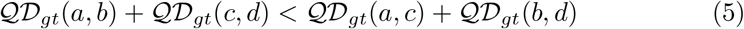

Thus, from equations 4 and 5, the following four-point condition holds for any arbitrary four taxa, and this completes the proof.

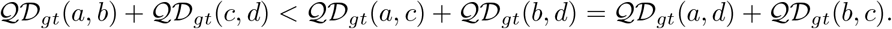

#### Theorem 2.

*The quartet distance is additive on the true species tree, and therefore STEQ is a statistically consistent method under the MSC model*.

*Proof*. The proof is provided in Supplementary Material (Section 1).

### 2.4 Normalized quartet distance computation

The standard quartet distance metric introduced in Section 2.2 quantifies the evolutionary dissimilarity between two taxa *x* and *y* based on the number of quartets in which they appear on opposite sides. While this method is statistically sound and efficient, it can disproportionately inflate distances for some pairs of taxa especially when an internal nodes of the gene tree define tripartitions with a very large third partition |*Z*|. In the original formula to calculate quartet distance between *x* and *y*, for the contribution of an internal node *u* with tripartition *X* |*Y* |*Z* (where *x* ∈ *X* and *y* ∈ *Y*), the terms (|*X* | *−*1) |*Z*| and (|*Y*| *−* 1) |*Z*| scale with |*Z*|. When |*Z*| is large, |*X*| and |*Y*| can be relatively very small in size, and can contribute disproportionately to the quartet distance calculation.

This scenario often occurs at shallow internal nodes, which are near the leaves of the tree and typically represent recent evolutionary divergences between closely related taxa. In this case, a small subset of closely related species may be grouped into small partitions, while the remaining majority of taxa may fall into a large third partition. For example, consider a gene tree *gt* comprising 100 taxa and an internal node *u* in *gt*, where species *w* and *x* form one partition *X*; species *y* and *z* form second partition *Y*, and the remaining 96 taxa are in the third partition *Z*. So *X* = {*w, x}, Y* = {*y, z*}, and *Z*={remaining 96 taxa}. Using the previously introduced unnormalized quartet distance formula,

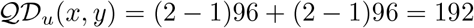

So a single internal node close to the leaves, although it represents very closely related taxa, can contribute as much as 192 units to the quartet distance between *x* and *y*, due to the large size of the unrelated group in |*Z*|. This exaggerated contribution misrepresents the actual evolutionary relationship between *x* and *y*, since it is primarily influenced by many unrelated taxa that should play only a minor role in their pairwise distance. With the growth of datasets containing a large number of taxa, the size of *Z* will also increase, making the effects more pronounced and underscoring the need for normalization to improve accuracy.

To address this problem, we introduce a normalized quartet distance that reduces the impact of large third partitions and places greater emphasis on the local topological structure. Rather than scaling by |*Z*|, the normalized quartet distance refines the contribution of each internal node *u* along the path between *x* and *y* as:

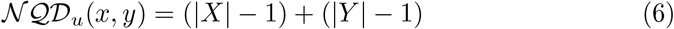

This eliminates the dependency on *Z*, which may include many distant or evolutionarily irrelevant taxa. The computation is analogous to the unnormalized case: for each pair (*x, y*), we identify internal nodes along their connecting path, compute their normalized contributions, sum them, and divide by 2 to avoid double-counting quartets (since each quartet maps to two nodes). The final distance matrix constructed from these values is then used in distancebased tree estimation algorithms such as FastME or BioNJ. Importantly, we have also proven that the normalized quartet distance retains additivity on the true species tree and is therefore statistically consistent under the multi-species coalescent (MSC) model (see Supplementary Section 2). So we now have the following Lemma:

#### Lemma 1.

*Let gt be an unrooted gene tree on a taxon set* 𝒳 *and u be an internal node of gt that induces a tripartition X*| *Y* |*Z of the taxon set* 𝒳. *Let x, y* ∈ *𝒳 be two distinct taxa such that x*∈ *X and y*∈ *Y*. *Then the normalized quartet distance between x and y associated with the node u is the total number of taxa in X and Y*, *excluding x and y themselves*.

## 3 Experimental studies

### 3.1 Datasets

#### Simulated datasets

We evaluated STEQ using large-scale simulated datasets previously analyzed in ASTRAL-II [23] and wQFM-TREE [26] studies. In these datasets, species trees were simulated according to the Yule process by varying the number of taxa (200, 500, and 1000 taxa), maximum tree length (500K, 2M and 10M generations) and speciation rate (1E − 6 and 1E − 7 per generation). Following ASTRAL-II, we organized our simulation study into two datasets. The first focuses on ILS by fixing number of taxa (200-taxon) and varying tree length and speciation rate. The second targets scalability by varying number of taxa (200, 500, and 1000) while fixing tree length (2M generations) and speciation rate (1E − 6). In both cases, we varied the number of genes (50, 200, and 1000). We analyzed all 50 replicates, except for the 1000-taxon condition where we ran STEQ on 20 replicates to match the original wQFM-TREE study. Since STEQ can’t handle polytomies, we resolved polytomies using the script from wQFM-TREE study (this preprocessing has no notable impact on the species tree accuracy [26]). We then applied STEQ on the resolved estimated gene trees. In addition, we evaluated all three methods on the 48-taxon avian and 37-taxon mammalian datasets from [21]. Supplementary Table S1 summarizes the parameter settings of all the simulated datasets.

#### Empirical datasets

We re-analyzed the green plant phylogenomic dataset from One Thousand Plant Transcriptomes Initiative (1KP, 2019) [1] containing 410 gene family trees and 1178 taxa. Moreover, we applied STEQ to the extended avian dataset [34] containing 363 species and 63,430 intergenic loci. For the empirical datasets, we also resolved the polytomy of the gene trees.

### 3.2 Methods and Measurements

We evaluated STEQ against the leading quartet based species tree estimation methods ASTRAL-III (v.5.7.8) [37], and wQFM-TREE [26]. On simulated datasets, the inferred trees were compared to the model species tree using normalized Robinson-Foulds (RF) distance [30]. For biological datasets, the estimated species trees were assessed qualitatively against well-established evolutionary relationships. We used Wilcoxon signed-rank test with *α* = 0.05 to measure the statistical significance of the differences between two methods.

## 4 Results and Discussion

We conducted two experiments on simulated datasets to evaluate: (i) the performance of STEQ under gene-tree discordance and (ii) the scalability as the number of taxa increases in large datasets. We also present the results on two biological datasets. All experiments have been performed on a Linux machine with a Intel® Core− i7 3.80 GHz processor and 64 GB of RAM.

### 4.1 Experiment 1: Performance of STEQ under Gene Tree Discordance

#### Results on 48-taxon dataset

Fig. 2(a) summarizes the accuracy of STEQ, ASTRAL-III, and wQFM-TREE on 48-taxon dataset using estimated gene trees from 500 bp sequences. Across the ILS-level model conditions, STEQ achieves the highest accuracy. It yields statistically significant improvements over ASTRAL-III in 2 of the 3 conditions, and over wQFM-TREE in 1 condition. With the increase of number of genes, STEQ remains consistently more accurate than ASTRAL-III across all settings, with statistically significant differences (*p <* 0.05) in 4 of the 5 conditions. Compared with wQFM-TREE, STEQ performs better in 3 of 5 conditions, achieving statistical significance in 2 of those 3.

**Fig. 2.**
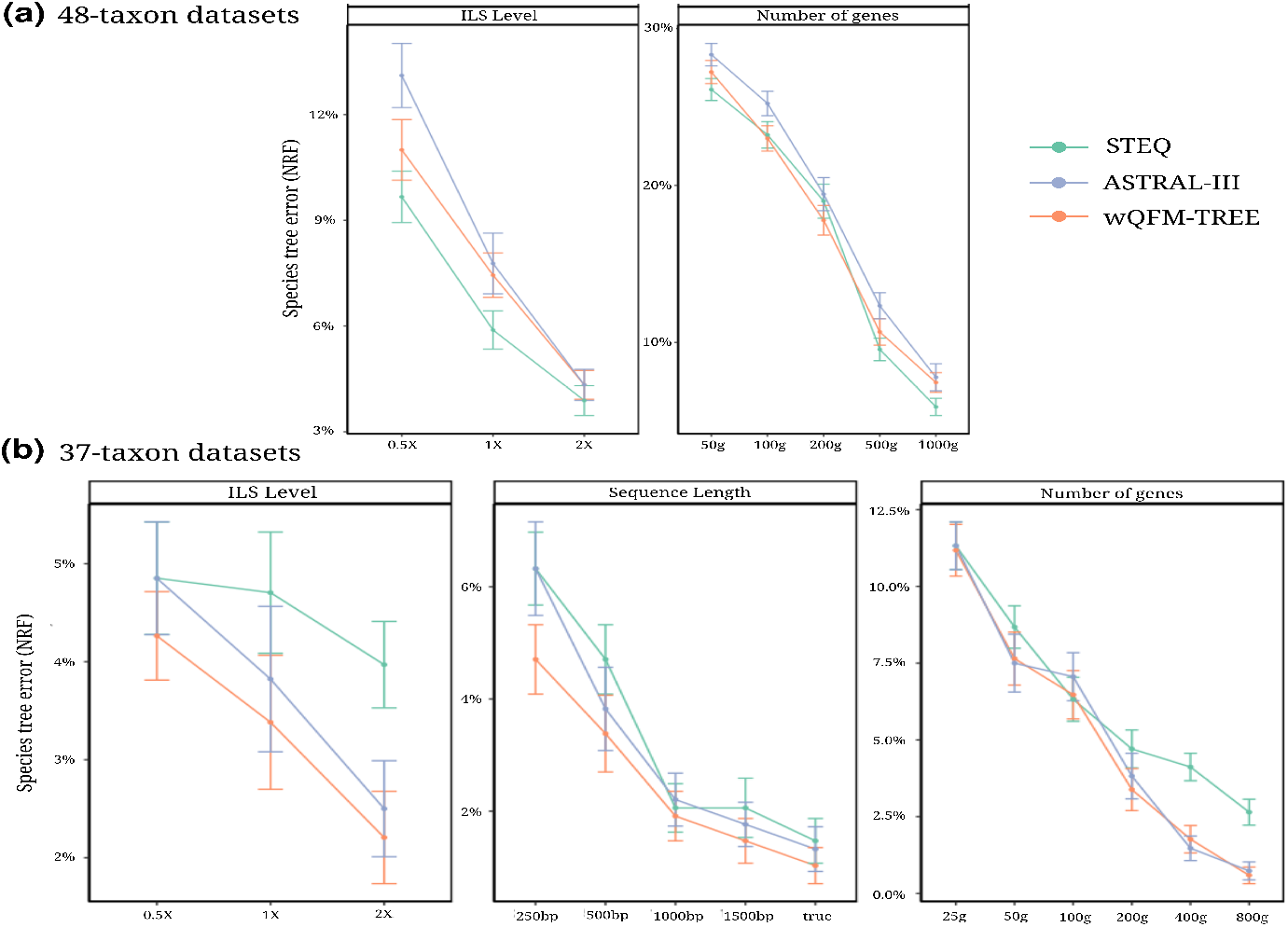
Average RF rates of STEQ, ASTRAL-III, and wQFM-TREE over 20 replicates (a) on 48-taxon dataset for estimated gene trees (500 bp sequence length) and (b) on 37-taxon dataset varying ILS level, number of genes and sequence length.

#### Results on 37-taxon dataset

Fig. 2(b) compares STEQ, ASTRAL-III, and wQFM-TREE on 37-taxon dataset across ILS levels, sequence lengths, and numbers of genes. STEQ remains competitive across all settings. Under low ILS (0.5X), STEQ matches the accuracy of ASTRAL-III. Considering both estimated and true gene trees (5 model conditions in total), STEQ and ASTRAL-III perform comparably, with no statistically significant differences (*p <* 0.05). STEQ outperforms ASTRAL-III for 1000 bp sequences and ties it for 250 bp sequences. Relative to wQFM-TREE, STEQ was outperformed with statistical significance in only 1 of 5 model conditions considering both estimated and true gene trees. Across gene counts, STEQ achieves the best performance with 100 genes and is also better than ASTRAL-III for 25 genes, although these differences are not statistically significant.

#### Results on 200-taxon dataset

Fig. 3(a) compares STEQ, ASTRAL-III, and wQFM-TREE on 200-taxon dataset varying speciation rates, tree lengths, and numbers of genes. For all three methods, accuracy improves as tree length and gene count increase. For short trees (500K generations), ASTRAL-III and wQFM-TREE slightly outperform STEQ overall. However, STEQ achieves the best performance in the 1000-gene setting under speciation rate 1E − 7, and it also improves upon wQFM-TREE under speciation rate 1E − 6. For moderate to long trees (2M and 10M generations), wQFM-TREE attains the lowest error among the three methods. Nevertheless, under speciation rate 1E − 6, STEQ is more accurate than ASTRAL-III in 4 out of 6 model conditions, although these differences are not statistically significant. Under speciation rate 1E − 7 with moderate to long trees, STEQ remains competitive.

**Fig. 3.**
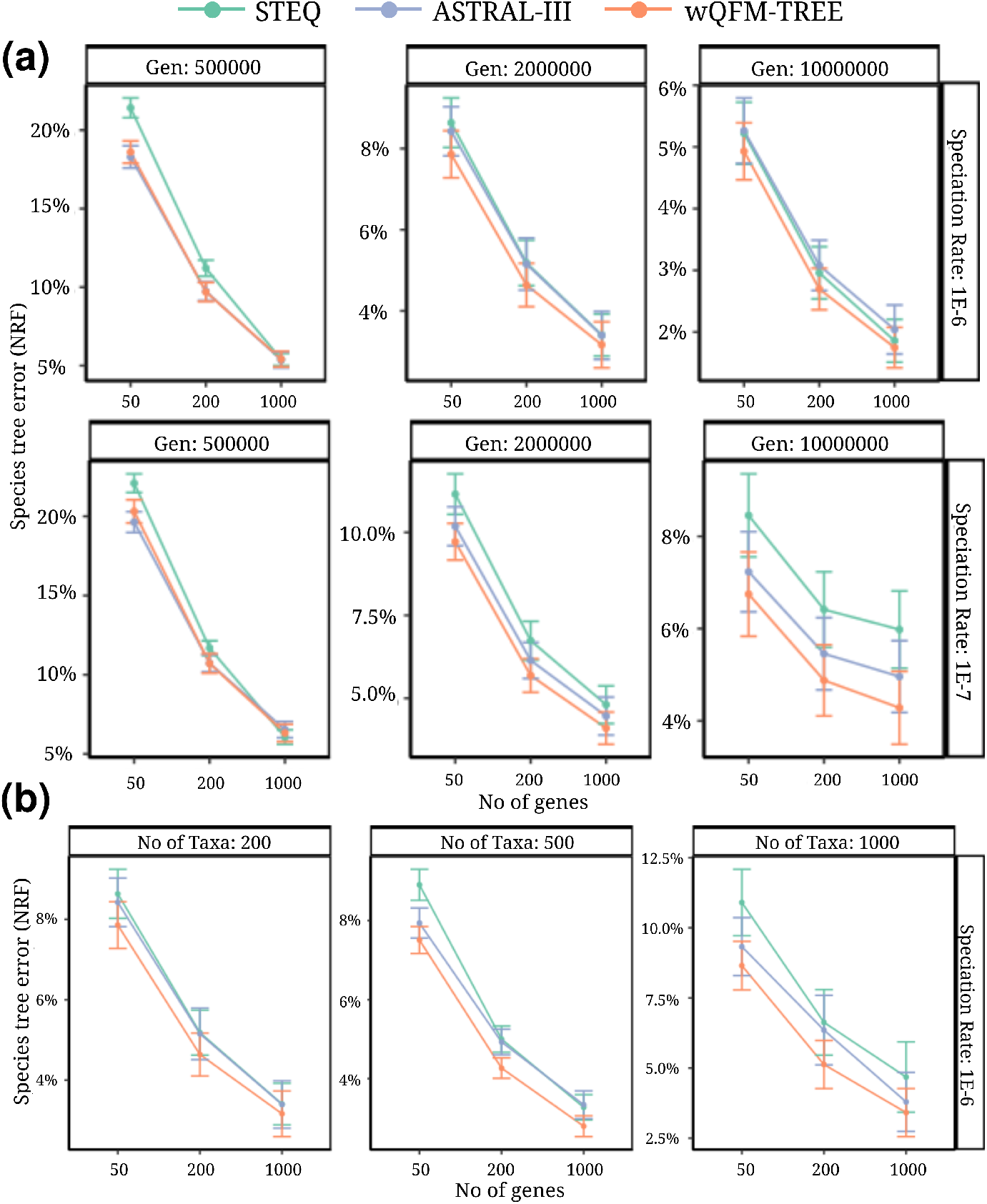
Average RF rates of three methods on datasets with (a) 200 taxa varying tree lengths (box column) and number of genes (x-axis) over 50 replicates, and with (b) 200, 500, and 1000 taxa varying number of genes with fixed tree length (2M generations) and fixed speciation rate (1E−6).

### 4.2 Experiment 2: Results on large datasets

Fig. 3(b) compares the three methods as the number of taxa increases (200, 500, and 1000), while fixing the tree length at 2M generations and speciation rate at 1E − 6. Consistent with Section 4.1, STEQ is more accurate than ASTRAL-III in 2 out of 3 model conditions for 200-taxon, although these differences are not statistically significant (*p* < 0.05). As the taxon count increases to 500 and 1000, STEQ remains competitive and achieves accuracy comparable to ASTRAL-III. ASTRAL-III achieves statistically significant improvement over STEQ in only 1 out of 6 model conditions on 500 and 1000 taxa. STEQ outperforms ASTRAL-III in one condition with 500 taxa and 1000 gene trees.

### 4.3 Results on empirical datasets

#### 1kp Dataset

We reanalyzed a plant transcriptome dataset comprising 1,178 species and 410 gene trees. The species tree inferred by STEQ is shown in Fig. 4(a). STEQ recovers all major clades, and the relationships among these clades are highly consistent with results from established methods. For reference, the species tree inferred by ASTRAL-III and wQFM-TREE are provided in the Fig. 4(a) and Supplementary Fig. S2 respectively. A detailed discussion of key phylogenetic relationships is included in the Supplementary Material (Section 4.1).

**Fig. 4.**
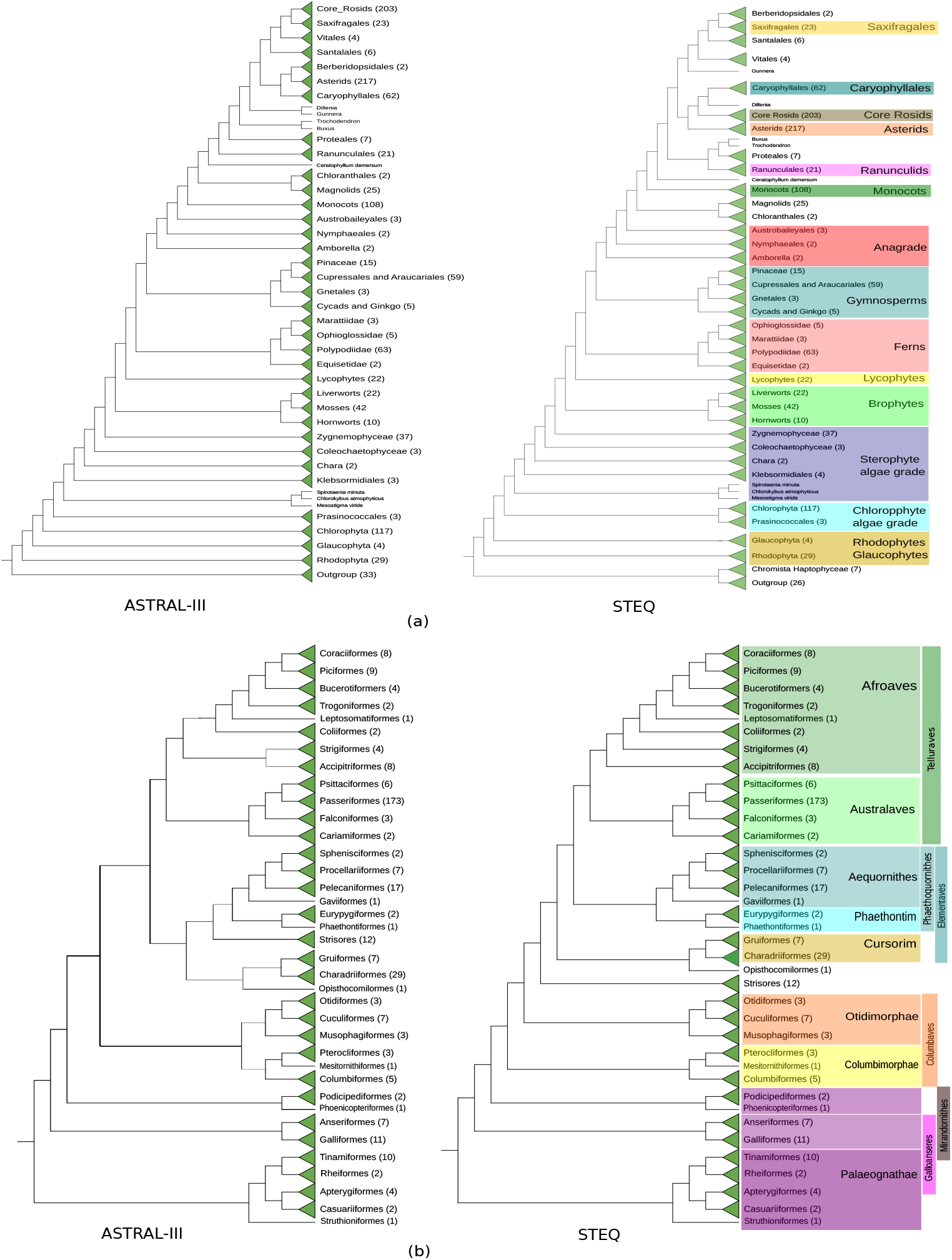
ASTRAL-III and STEQ estimated species trees using (a) plant transcriptome dataset comprising 1,178 species and 410 gene trees and (b) extended avian dataset comprising 363 species and 63430 gene trees (intergenic loci).

#### Extended Avian Dataset

We analyzed an extended avian dataset [34] comprising 363 species and 63430 gene trees (intergenic loci). STEQ recovered the three major avian groups—Palaeognathae (ratites and tinamous), Galloanseres (landfowl and waterfowl), and Neoaves. Within Neoaves, STEQ correctly reconstructed all the major clades reported in [34], including Mirandornithes, Columbimorphae, Otidimorphae, Strisores, Opisthocomiformes, Cursorimorphae, Aequornithes, Phaethontimorphae, and Telluraves. Most inter-clade relationships inferred by STEQ are consistent with those supported by established methods (see Fig. 4(b)). The species tree inferred by wQFM-TREE is highly similar to the STEQ tree (see Supplementary Fig. S3). Key phylogenetic relationships are discussed in details in the Supplementary Material (Section 4.2).

### 4.4 Running Time

STEQ proceeds in two phases: (i) computing the *n* × *n* distance matrix ℳ, and (ii) inferring a species tree from ℳ using a distance-based method.

#### Phase I: distance matrix computation

For each unordered taxon pair {*x, y*}, STEQ scans the *k* gene trees and computes 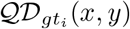 only for those *gt*_*i*_ that contain both *x* and *y*. For a fixed gene tree, computing 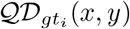 traverses exactly the internal nodes *u* ∈ *U*_*xy*_ and accumulates the contribution at each visited node (Eqn. 2). A standard precomputation with bottom-up dynamic programming enables *O*(1) lookup of the contribution at any visited internal node. Hence, the per-tree cost for a pair {*x, y*} is linear in the number of internal nodes on the unique *x*–*y* path.

##### Theorem 3.

*Let T be a rooted perfect binary tree with n* = 2^*d*^ *leaves. For two distinct leaves x and y*,

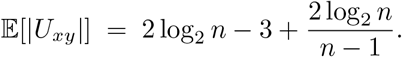

*Proof*. Proof is provided in Supplementary Material (Section 3).

When gene trees are reasonably balanced, Theorem 3 implies that the expected number of visited internal nodes per pair is *Θ*(log *n*), and therefore the expected cost to compute 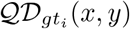 for a single gene tree is *O*(log *n*). Since there are 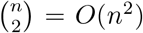 taxon pairs and distances are aggregated across *k* gene trees, the expected total time to compute the matrix ℳ is *O*(*kn*^2^ log *n*). In the worst case of highly unbalanced (caterpillar) gene trees, *U*_*xy*_ can contain *Θ*(*n*) internal nodes, and the matrix computation may require *O*(*kn*^3^).

#### Phase II: species tree reconstruction

Given ℳ, STEQ reconstructs a tree using FastME when ℳ has no missing entries and BioNJ otherwise. This phase depends only on *n* (not on *k*) and runs in polynomial time in *n* (typically between *O*(*n*^2^) and *O*(*n*^3^), depending on the implementation).

#### Overall complexity

Thus, in the common case *k* ≫ *n* in phylogenomic datasets, the total running time is dominated by Phase I and is *O*(*kn*^2^ log *n*) for reasonably balanced gene trees, and *O*(*kn*^3^) in the worst case.

#### Running time on simulated datasets

We assessed the computational efficiency of STEQ, ASTRAL-III, and wQFM-TREE on simulated datasets spanning 37–1000 taxa and 50–1000 gene trees (Supplementary Table S2). Across all settings, STEQ was substantially faster. On the 200-taxon dataset with 1000 genes, STEQ completed within 30 seconds, whereas ASTRAL-III and wQFM-TREE required around 4–6 minutes. For 500 taxa, STEQ’s maximum running time was about 4 minutes, compared with 25–40 minutes for ASTRAL-III and wQFM-TREE. On the largest 1000-taxon dataset, STEQ finished in under 20 minutes, while the other two methods required about 2–3 hours.

#### Running time on empirical datasets

We also compared runtimes on two empirical datasets. On the plant transcriptome dataset (1,178 taxa; 410 genes), STEQ finished in ~7 minutes, whereas ASTRAL required ~1 hour and wQFM-TREE nearly 3 hours. On the extended avian dataset (363 species; 63,430 gene trees), ASTRAL took almost 1 day and wQFM-TREE about 2.5 days, while STEQ completed within only 3 hours.

## 5 Conclusions

In this paper, we introduced STEQ, a scalable distance-based method for species tree estimation from gene trees. STEQ computes a quartet-based distance matrix, bridging classic distance-based approaches that rely on internal nodes or edges and coalescent-based summary methods that maximize quartet agreement with input gene trees. Despite its simple formulation, STEQ scales to datasets with thousands of taxa and gene trees enabling significantly faster inference while maintaining competitive accuracy. We also prove that STEQ is statistically consistent under the multi-species coalescent (MSC) model. Across extensive evaluation on simulated and empirical datasets, STEQ matches the accuracy of established quartet-based methods, including ASTRAL and wQFM-TREE. On empirical datasets it recovers nearly all major clades and key evolutionary relationships supported by other widely used approaches. Compared with wQFM and wQFM-TREE, which often face scalability bottlenecks as taxon counts grow, STEQ substantially reduces running time without sacrificing accuracy. Our current implementation is single-core and our runtime comparisons therefore reflect single-core executions. Future work includes developing a multiparallel implementation of STEQ and extending the framework to triplet-based distance metrics and multi-copy gene trees.

## Supporting information

Suppementary Materials with Proofs, Tables, and Figures

