## Supplementary material for "STEQ: A statistically consistent quartet distance based species tree estimation method": Suppementary Materials with Proofs, Tables, and Figures

This supplementary material presents additional theorems and proofs on quartet distance, normalized quartet distance, and running time analysis. Here we also present detailed discussions on key phylogenetic relationships among different clades for empirical datasets, and additional figures and tables.

### 1 Statistical Consistency with Quartet Distance

**Theorem 1.** *The quartet distance is additive on the true species tree, and therefore STEQ is a statistically consistent method under the MSC model.*

*Proof.* Let $ST$ be the true species tree on $n$ taxa, and fix four arbitrary taxa $a, b, c, d \in L(ST)$. Without loss of generality, assume that the quartet induced by $ST$ on $\{a, b, c, d\}$ is $q = ab|cd$.

For each gene tree $gt_i \in \mathcal{G}$, define the three values corresponding to the three quartet topologies:

$$V_{gt_i}(ab|cd) = \mathcal{QD}_{gt_i}(a, b) + \mathcal{QD}_{gt_i}(c, d),$$
$$V_{gt_i}(ac|bd) = \mathcal{QD}_{gt_i}(a, c) + \mathcal{QD}_{gt_i}(b, d),$$
$$V_{gt_i}(ad|bc) = \mathcal{QD}_{gt_i}(a, d) + \mathcal{QD}_{gt_i}(b, c).$$

We also define their sums across the $k$ gene trees:

$$V(ab|cd) := \sum_{i=1}^{k} V_{gt_i}(ab|cd), \quad V(ac|bd) := \sum_{i=1}^{k} V_{gt_i}(ac|bd), \quad V(ad|bc) := \sum_{i=1}^{k} V_{gt_i}(ad|bc).$$

To prove additivity on $ST$, it suffices to establish the four-point condition for the quartet $\{a, b, c, d\}$, namely

$$V(ab|cd) \; < \; V(ac|bd) \; = \; V(ad|bc), \tag{1}$$

with probability tending to 1 as $k \to \infty$ (equivalently, in expectation as $k \to \infty$).

**Step 1: Per-locus difference random variables.** Let us define for each locus $i$ the differences

$$D_i^{(1)} := V_{gt_i}(ac|bd) - V_{gt_i}(ab|cd), \; D_i^{(2)} := V_{gt_i}(ad|bc) - V_{gt_i}(ab|cd), \; D_i^{(3)} := V_{gt_i}(ac|bd) - V_{gt_i}(ad|bc).$$

Then

$$V(ac|bd) - V(ab|cd) = \sum_{i=1}^{k} D_i^{(1)}, \; V(ad|bc) - V(ab|cd) = \sum_{i=1}^{k} D_i^{(2)}, \; V(ac|bd) - V(ad|bc) = \sum_{i=1}^{k} D_i^{(3)}.$$

Thus, to prove (1), it is enough to show that with probability $\to 1$ as $k \to \infty$,

$$\sum_{i=1}^{k} D_i^{(1)} > 0, \qquad \sum_{i=1}^{k} D_i^{(2)} > 0, \qquad \sum_{i=1}^{k} D_i^{(3)} = 0.$$

**Step 2: Values of $D_i^{(1)}, D_i^{(2)}, D_i^{(3)}$ depend only on the induced gene-tree quartet.** Let us consider the restriction of $gt_i$ to the four taxa $\{a, b, c, d\}$. There are three possible quartet topologies.

– If $gt_i$ induces $ab|cd$, then by Theorem 1 in the main paper,

$$V_{gt_i}(ab|cd) < V_{gt_i}(ac|bd) = V_{gt_i}(ad|bc).$$

Hence $D_i^{(1)} > 0$, $D_i^{(2)} > 0$, and $D_i^{(3)} = 0$. Since $\mathcal{QD}_{gt_i}(\cdot, \cdot)$ are integer counts, these imply $D_i^{(1)} \geq 1$ and $D_i^{(2)} \geq 1$.

– If $gt_i$ induces $ac|bd$, then by Theorem 1 in the main paper,

$$V_{gt_i}(ac|bd) < V_{gt_i}(ab|cd) = V_{gt_i}(ad|bc).$$

Hence $D_i^{(1)} < 0$, $D_i^{(2)} = 0$, and $D_i^{(3)} < 0$, and by integrality $D_i^{(1)} \leq -1$ and $D_i^{(3)} \leq -1$.

– If $gt_i$ induces $ad|bc$, then similarly

$$V_{gt_i}(ad|bc) < V_{gt_i}(ab|cd) = V_{gt_i}(ac|bd).$$

Hence $D_i^{(2)} < 0$, $D_i^{(1)} = 0$, and $D_i^{(3)} > 0$, and by integrality $D_i^{(2)} \leq -1$ and $D_i^{(3)} \geq 1$.

**Step 3: Positive expected drift under the MSC model.** Under the MSC model on the 4-taxon species quartet $ab|cd$, the matching gene-tree quartet topology $ab|cd$ occurs with probability $p > 1/3$, and each discordant topology occurs with probability $(1-p)/2$. Therefore,

$$\begin{aligned}\mathbb{E}[D_i^{(1)}] &\geq 1 \cdot \Pr(gt_i|_{\{a,b,c,d\}} = ab|cd) + (-1) \cdot \Pr(gt_i|_{\{a,b,c,d\}} = ac|bd) + 0 \cdot \Pr(gt_i|_{\{a,b,c,d\}} = ad|bc) \\ &= p - \frac{1-p}{2} = \frac{3p-1}{2} > 0,\end{aligned}$$

and similarly $\mathbb{E}[D_i^{(2)}] = \frac{3p-1}{2} > 0$.

To show the equality $V(ac|bd) = V(ad|bc)$, note that under the same MSC model the two discordant topologies $ac|bd$ and $ad|bc$ occur with equal probability $(1-p)/2$, and are symmetric under swapping the labels $c$ and $d$. This symmetry swaps $V_{gt_i}(ac|bd)$ and $V_{gt_i}(ad|bc)$, and hence maps $D_i^{(3)}$ to $-D_i^{(3)}$ while preserving its probability. Therefore, $\mathbb{E}[D_i^{(3)}] = 0$.

**Step 4: Concluding the four-point condition by the law of large numbers.** Assuming loci are sampled independently, $\{D_i^{(1)}\}_{i=1}^k$, $\{D_i^{(2)}\}_{i=1}^k$, and $\{D_i^{(3)}\}_{i=1}^k$ are i.i.d. with finite expectation. By the strong law of large numbers (here a.s. means "almost surely"),

$$\frac{1}{k}\sum_{i=1}^{k} D_i^{(1)} \xrightarrow[k\to\infty]{a.s.} \mathbb{E}[D_i^{(1)}] > 0, \quad \frac{1}{k}\sum_{i=1}^{k} D_i^{(2)} \xrightarrow[k\to\infty]{a.s.} \mathbb{E}[D_i^{(2)}] > 0, \quad \frac{1}{k}\sum_{i=1}^{k} D_i^{(3)} \xrightarrow[k\to\infty]{a.s.} \mathbb{E}[D_i^{(3)}] = 0.$$

Hence, with probability tending to 1 as $k \to \infty$, we have

$$\sum_{i=1}^{k} D_i^{(1)} > 0, \qquad \sum_{i=1}^{k} D_i^{(2)} > 0, \qquad \sum_{i=1}^{k} D_i^{(3)} = 0,$$

which implies

$$V(ab|cd) < V(ac|bd) = V(ad|bc) \text{ in the limit.}$$

If gene trees may be incomplete, the same reasoning applies after restricting to those loci for which all four taxa $a, b, c, d$ are present. Thus we obtain,

$$\mathcal{QD}(a,b) + \mathcal{QD}(c,d) < \mathcal{QD}(a,c) + \mathcal{QD}(b,d) = \mathcal{QD}(a,d) + \mathcal{QD}(b,c).$$

Therefore, the four-point condition holds for any arbitrary four taxa given a sufficiently large number of true gene trees. Therefore, as Saitou and Nei showed (4), the NJ method, run on the distance matrix constructed using the quartet distance, will return the true species tree. This completes the proof.

### 2 Normalized Quartet Distance

**Theorem 2.** *The normalized quartet distance is additive on a gene tree.*

*Proof.* To prove that normalized quartet distance is additive, it is sufficient to prove that the four-point condition holds for any arbitrary set of four species. Let $gt$ be a true gene tree for a particular gene sampled from $n$ species. Consider four arbitrary taxa $a, b, c, d$ in $gt$. Without loss of generality, assume that $a$ and $b$ are more closely related to each other than they are to $c$ and $d$ in $gt$. That means $q = ab|cd$ is the quartet induced by these four taxa in $gt$. Let $q$ maps to two internal nodes $u$ and $u'$ (see Figure S1).

The normalized distance between $\mathcal{NQD}_{gt}(a,c)$ between $a$ and $c$ can be calculated as follows:

$\mathcal{NQD}_{gt}(a,c) = \sum_{u \in U_{ac}} \mathcal{NQD}_u(a,c)$

$U_{ac}$ contains all the internal nodes on the path from $a$ to $c$ in $gt$. We divide $U_{ac}$ into 5 subsets:

i) $U_{ac}^1 = U_{au}$ (internal nodes from $a$ to $u$ excluding $u$),

ii) $U_{ac}^2 = \{u\}$,

iii) $U_{ac}^3 = U_{uu'}$ (internal nodes from $u$ to $u'$ excluding $u$ and $u'$),

iv) $U_{ac}^4 = \{u'\}$, and

v) $U_{ac}^5 = U_{u'c}$ (internal nodes from $u'$ to $c$ excluding $u'$)

Thus, $U_{ac} = U_{ac}^1 \cup U_{ac}^2 \cup U_{ac}^3 \cup U_{ac}^4 \cup U_{ac}^5$.

Let the number of taxa descending from the internal nodes on the path from $a$ to $u$ (excluding $u$) is $A$, and $B$ be the number of taxa descending from the internal nodes on the path from $b$ to $u$ (excluding $u$). Similarly, we can define $C$ and $D$ with respect to the other mapping node $u'$.

We now define the following distances for computational convenience.

$\sum_{u \in U_{au}} \mathcal{NQD}_u(a,c) = \sum_{u \in U_{au}} \mathcal{NQD}_u(a,b) = \sum_{u \in U_{au}} \mathcal{NQD}_u(a,d) = \mathcal{AU}$

$\sum_{u \in U_{cu'}} \mathcal{NQD}_u(a,c) = \sum_{u \in U_{cu'}} \mathcal{NQD}_u(b,c) = \sum_{u \in U_{cu'}} \mathcal{NQD}_u(c,d) = \mathcal{CU}'$

$\sum_{u \in U_{uu'}} \mathcal{NQD}_u(a,c) = \sum_{u \in U_{uu'}} \mathcal{NQD}_u(a,d) = \sum_{u \in U_{uu'}} \mathcal{NQD}_u(b,c) = \sum_{u \in U_{uu'}} \mathcal{NQD}_u(b,d) = \mathcal{UU}'$

Similarly, we can define $\mathcal{BU}$ and $\mathcal{DU}'$.

Now,

$$\begin{aligned}
\mathcal{NQD}_{gt}(a,c) &= \frac{1}{2} \sum_{u \in U_{ac}} \mathcal{NQD}_u(a,c) \\
&= \frac{1}{2} \sum_{i=1}^{5} \sum_{u \in U_{ac}^i} \mathcal{NQD}_u(a,c) \\
&= \frac{1}{2} \big( \mathcal{AU} + \mathcal{NQD}_u(a,c) + \mathcal{UU}' + \mathcal{NQD}_{u'}(a,c) + \mathcal{CU}' \big)
\end{aligned}$$

Here, according to Equation 6 in the main paper,

$\mathcal{NQD}_u(a,c) = (A-1) + (n-A-B-1)$, and
$\mathcal{NQD}_{u'}(a,c) = (C-1) + (n-C-D-1)$. Therefore,

$$\mathcal{NQD}_{gt}(a,c) = \frac{1}{2}\{\mathcal{AU} + (A-1) + (n-A-B-1) + \mathcal{UU'} + (C-1) + (n-C-D-1) + \mathcal{CU'}\}$$

Similarly,

$$\mathcal{NQD}_{gt}(b,d) = \frac{1}{2}\{\mathcal{BU} + (B-1) + (n-A-B-1) + \mathcal{UU'} + (D-1) + (n-C-D-1) + \mathcal{DU'}\}$$

$$\mathcal{NQD}_{gt}(a,d) = \frac{1}{2}\{\mathcal{AU} + (A-1) + (n-A-B-1) + \mathcal{UU'} + (D-1) + (n-C-D-1) + \mathcal{DU'}\}$$

$$\mathcal{NQD}_{gt}(b,c) = \frac{1}{2}\{\mathcal{BU} + (B-1) + (n-A-B-1) + \mathcal{UU'} + (C-1) + (n-C-D-1) + \mathcal{CU'}\}$$

$$\mathcal{NQD}_{gt}(a,b) = \frac{1}{2}\{\mathcal{AU} + (A-1) + (B-1) + \mathcal{BU}\}$$

$$\mathcal{NQD}_{gt}(c,d) = \frac{1}{2}\{\mathcal{CU'} + (C-1) + (D-1) + \mathcal{DU'}\}$$

Now,

$$\begin{aligned}
&\mathcal{NQD}_{gt}(a,c) + \mathcal{NQD}_{gt}(b,d) \\
&= \frac{1}{2}\{\mathcal{AU} + \mathcal{BU} + \mathcal{CU'} + \mathcal{DU'} + 2\mathcal{UU'} + (A-1) + (B-1) + (C-1) + (D-1) \\
&+ 2(n-A-B-1) + 2(n-C-D-1)\} \\
&= \frac{1}{2}\{\mathcal{AU} + \mathcal{BU} + \mathcal{CU'} + \mathcal{DU'} + 2\mathcal{UU'} + (A-1) + (B-1) + (C-1) + (D-1)\} \\
&+ \{2n - (A+B+C+D) - 2\}
\end{aligned}$$

Similarly, it is easy to see that

$$\begin{aligned}
&\mathcal{NQD}_{gt}(a,d) + \mathcal{NQD}_{gt}(b,c) \\
&= \frac{1}{2}\{\mathcal{AU} + \mathcal{BU} + \mathcal{CU'} + \mathcal{DU'} + 2\mathcal{UU'} + (A-1) + (B-1) + (C-1) + (D-1)\} \\
&+ \{2n - (A+B+C+D) - 2\}
\end{aligned}$$

Therefore,

$$\mathcal{NQD}_{gt}(a,c) + \mathcal{NQD}_{gt}(b,d) = \mathcal{NQD}_{gt}(a,d) + \mathcal{NQD}_{gt}(b,c) \tag{2}$$

We now compute the value of $\mathcal{NQD}_{gt}(a,b) + \mathcal{NQD}_{gt}(c,d)$.

$$\begin{aligned}
&\mathcal{NQD}_{gt}(a,b) + \mathcal{NQD}_{gt}(c,d) \\
&= \frac{1}{2}\{\mathcal{AU} + \mathcal{BU} + \mathcal{CU'} + \mathcal{DU'} + 2\mathcal{UU'} + (A-1) + (B-1) + (C-1) + (D-1)\}
\end{aligned}$$

Note that $A+B+C+D \leq n$. So,

$$2n - (A+B+C+D) - 2 \geq n-2.$$

Since we are considering unrooted tree of four taxa, $n \geq 4$ or $n-2 \geq 2$ and so,

$$2n - (A+B+C+D) - 2 \geq 2.$$

Therefore,

$$\mathcal{NQD}_{gt}(a,b) + \mathcal{NQD}_{gt}(c,d) < \mathcal{NQD}_{gt}(a,c) + \mathcal{NQD}_{gt}(b,d) \tag{3}$$

Thus, from equations 2 and 3, the following four-point condition holds for any arbitrary four taxa, and this completes the proof.

$$\mathcal{NQD}_{gt}(a,b) + \mathcal{NQD}_{gt}(c,d) < \mathcal{NQD}_{gt}(a,c) + \mathcal{NQD}_{gt}(b,d) = \mathcal{NQD}_{gt}(a,d) + \mathcal{NQD}_{gt}(b,c).$$

**Theorem 3.** *The normalized quartet distance is additive on the true species tree, and therefore STEQ with normalized quartet distance is a statistically consistent method under the MSC model.*

*Proof.* Let us fix any four distinct taxa $a,b,c,d \in \mathcal{X}$, and let the quartet induced by the true species tree $ST$ on $\{a,b,c,d\}$ be $ab|cd$. For each locus $i$, let us define

$$V_{gt_i}(ab|cd) = \mathcal{NQD}_{gt_i}(a,b) + \mathcal{NQD}_{gt_i}(c,d)$$

$$V_{gt_i}(ac|bd) = \mathcal{NQD}_{gt_i}(a,c) + \mathcal{NQD}_{gt_i}(b,d)$$

$$V_{gt_i}(ad|bc) = \mathcal{NQD}_{gt_i}(a,d) + \mathcal{NQD}_{gt_i}(b,c)$$

By Theorem 2, for each $i$ the unique minimum among these three values corresponds exactly to the quartet topology induced by $gt_i$ on $\{a,b,c,d\}$ (with the other two equal).

Now following the species-tree additivity proof in the main paper and replacing $\mathcal{QD}$ by $\mathcal{NQD}$: let us define the per-locus differences $D_i^{(1)} := V_{gt_i}(ac|bd) - V_{gt_i}(ab|cd)$, $D_i^{(2)} := V_{gt_i}(ad|bc) - V_{gt_i}(ab|cd)$, and $D_i^{(3)} := V_{gt_i}(ac|bd) - V_{gt_i}(ad|bc)$. Under the MSC on the species quartet $ab|cd$, the matching gene-quartet occurs with probability $p > 1/3$ and each discordant quartet with probability $(1-p)/2$, implying $\mathbb{E}[D_i^{(1)}] > 0$ and $\mathbb{E}[D_i^{(2)}] > 0$, while symmetry of the two discordant topologies gives $\mathbb{E}[D_i^{(3)}] = 0$. Assuming loci are independent, the strong law of large numbers yields

$$\sum_{i=1}^{k} V_{gt_i}(ab|cd) \; < \; \sum_{i=1}^{k} V_{gt_i}(ac|bd) \; = \; \sum_{i=1}^{k} V_{gt_i}(ad|bc) \quad \text{with probability } \to 1 \text{ as } k \to \infty,$$

equivalently,

$$\mathcal{NQD}(a,b) + \mathcal{NQD}(c,d) \; < \; \mathcal{NQD}(a,c) + \mathcal{NQD}(b,d) \; = \; \mathcal{NQD}(a,d) + \mathcal{NQD}(b,c).$$

Thus $\mathcal{NQD}$ satisfies the four-point condition on $ST$ (in the large-locus limit), so it is additive on the true species tree and STEQ with $\mathcal{NQD}$ is statistically consistent under the MSC.

### 3 Running Time Analysis

**Theorem 4.** *Let $T$ be a rooted perfect binary tree with $n = 2^d$ leaves. For two distinct leaves $x$ and $y$, the average number of internal nodes on the unique $x$–$y$ path is given by,*

$$\mathbb{E}[|U_{xy}|] \; = \; 2\log_2 n - 3 + \frac{2\log_2 n}{n-1}.$$

*Proof.* All leaves have depth $d$. If the lowest common ancestor of $x$ and $y$ denoted by $\mathrm{LCA}(x, y)$ has depth $t \in \{0, \dots, d-1\}$, then the unique $x$–$y$ path has $2(d-t)$ edges, so it contains $2(d-t)+1$ vertices, of which two are leaves; hence the number of internal vertices on the path is

$$|U_{xy}| = 2(d-t) - 1. \tag{4}$$

At depth $t$ there are $2^t$ nodes, and for each such node $v$ its two child subtrees contain $2^{d-t-1}$ leaves each. Thus the number of unordered leaf pairs whose LCA is exactly $v$ equals $2^{d-t-1} \cdot 2^{d-t-1} = 2^{2(d-t-1)}$, and therefore the total number of pairs with LCA depth $t$ is

$$N_t = 2^t \cdot 2^{2(d-t-1)} = 2^{2d-t-2}. \tag{5}$$

Summing (4) over all pairs and dividing by $\binom{n}{2} = \binom{2^d}{2} = 2^{d-1}(2^d - 1)$ yields

$$\mathbb{E}[|U_{xy}|] = \frac{\sum_{t=0}^{d-1} N_t \,(2(d-t)-1)}{\binom{2^d}{2}} = 2(d-1) + \frac{2d}{2^d - 1} - 1 = 2d - 3 + \frac{2d}{2^d - 1}.$$

Substituting $d = \log_2 n$ gives the stated expression.

### 4 Results on Empirical Datasets

#### 4.1 1kp Dataset

*Primary acquisition of the plastid* The relationship among Viridiplantae, Glaucophyta, and Rhodophyta is of fundamental evolutionary importance, as it relates to the origin of plastids—a defining event in the history of photosynthetic eukaryotes. In our analysis, STEQ recovered a sister relationship between Viridiplantae and Glaucophyta similar to ASTRAL-III and wQFM-TREE, which suggests that the ancestral red algae experienced the loss of flagella and peptidoglycan biosynthesis.

*Viridiplantae* STEQ successfully recovered the monophyletic relationship of Viridiplantae, with early-diverging Chlorophyta and Streptophyta forming distinct and consistent lineages—aligning well with previous phylogenomic studies. However, the placement of Prasinococcales is found to be unstable in 1 kp (2019). Unlike ASTRAL-III and wQFM-TREE, STEQ placed Prasinococcales as a sister to Chlorophyta.

*Diversification within Chlorophyta* In our analysis, STEQ recovered Trebouxiophyceae as sister to the clade containing Chlorophyceae and Ulvophyceae, consistent with the topology recovered by ASTRAL-III. Notably, STEQ successfully reconstructed the monophyletic relationships within Briopsidales (an order of green algae, in the class Ulvophyceae) and other Ulvophyceae. For both ASTRAL-III and wQFM-TREE, Briopsidales did not exhibit monophyly with other Ulvophyceae. Instead, it was positioned as a sister group to Pedinophyceae for wQFM-TREE and was positioned as a sister to Chlorophyceae for ASTRAL-III.

*Streptophyta* STEQ recovered the internal relationships of Streptophyta consistent with both ASTRAL-III and wQFM-TREE. Specifically, *Mesostigma*, *Spirotaenia minuta*, and *Chlorokybus* formed a clade that is sister to the remainder of Streptophyta, with a well-supported pattern of successive divergence involving Klebsormidiales, Charophyceae, Coleochaetophyceae, and Zygnematophyceae, relative to Embryophyta.

*Embryophyta* Although the relationships among Bryophytes—including mosses, liverworts, and hornworts—have been contentious, with some studies resolving them as a paraphyletic grade, STEQ recovered Bryophytes as a monophyletic group. This result is also consistent with both ASTRAL-III and wQFM-TREE, which rejected the hypothesis that liverworts are sister to all other extant land plant lineages.

*Vascular plants* STEQ correctly recovered Lycophytes as the sister group to ferns and seed plants, consistent with ASTRAL-III and wQFM-TREE. In the case of Marattiales, a group whose phylogenetic placement has been debated, STEQ's result agrees with ASTRAL-III, placing Marattiales as sister to Ophioglossidae. This contrasts with the placement inferred by wQFM-TREE, which positioned Marattiales as sister to Polypodiidae.

*Seed plants* STEQ correctly recovered Gymnosperms as the sister group to flowering plants, consistent with widely accepted evolutionary relationships. However, the placement of Gnetales within gymnosperms has been a subject of long-standing debate, giving rise to several competing hypotheses: "Gnecup", "Gnepine", and "Gnetifer". In our analysis, STEQ supported the "Gnetifer" hypothesis, placing Gnetales as sister to all other conifers (i.e., Araucariales, Cupressales, and Pinales), consistent with ASTRAL-III. This result contrasts with wQFM-TREE, which favored the "Gnepine" hypothesis, placing Gnetales as sister to Pinales. In our analysis, the relationships among early-diverging angiosperm lineages were consistent with those recovered by ASTRAL-III and wQFM-TREE. Specifically, Amborellales, Nymphaeales, and Austrobaileyales were placed as successive sisters to all other angiosperms, supporting a widely accepted model of basal angiosperm diversification. Furthermore, Chloranthales and Magnoliids were grouped as sister clades, and this pair was resolved as successive sister lineages to the remaining Mesangiospermae, which includes monocots, Ceratophyllum, and eudicots.

Several topological differences were observed among the eudicots when comparing STEQ with ASTRAL-III and wQFM-TREE. ASTRAL-III and wQFM-TREE recovered a monophyletic clade comprising Core Rosids, Saxifragales, Vitales, and Santalales, whereas STEQ resolved these lineages as a paraphyletic grade, suggesting more gradual divergence among early-diverging core eudicots. In contrast, STEQ placed *Buxus* and *Trochodendron* as sister taxa, consistent with ASTRAL-III, while wQFM-TREE grouped *Trochodendron* as sister to the clade consisting *Buxus* and core eudicots. Additionally, the placement of *Gunnera* and *Dillenia* in STEQ's estimated tree diverged from both ASTRAL-III and wQFM-TREE.

#### 4.2 Extended Avian Dataset

*Mirandornithes placed as sister to other Neoaves* The placement of Mirandornithes as the sister lineage to the remaining Neoaves was supported by all the methods: ASTRAL, concatenation, STEQ, and wQFM-TREE. Similar to concatenation (5) , STEQ and wQFM-TREE placed Mirandornithes and Columbimorphae as successive sister groups to the remaining neoavian clades. However, ASTRAL combined Columbimorphae with Otidimorphae to form the clade Columbave, which has also been reported previously, albeit with low bootstrap support (3). Within Otidimorphae, all methods resolved Otidiformes as the sister group to Cuculiformes.

*Waterbirds placed in a diverse clade deep in the lineage* STEQ and wQFM-TREE placed Phaethoquornithes (Aequornithes + Phaethontimorphae) as sister to landbirds in accordance with previous hypotheses (1), where ASTRAL placed Phaethoquornithes deep inside the diverse clade Elementaves (containing Aequornithes + Phaethontimorphae, Strisores, Opisthocomiformes and Cursorimorphae). STEQ and wQFM-TREE consistently grouped Charadriiformes and Gruiformes (forming Cursorimorphae), and placed the third orphan, Opisthocomiformes, as the sister to this group similar to ASTRAL. ASTRAL placed Strisores with Phaethoquornithes with moderate support, where STEQ and wQFM-TREE placed Strisores as a sister lineage with Opisthocomiformes + Cursorimorphae. Previous studies also failed to find unequivocal support for the relationship of Strisores, placing it as sister to Otidimorphae (1), Cursorimorphae (6) or Opisthocomiformes (2).

*Difficult placement of owls and hawks* Within Telluraves, all the methods agreed with the proposed split into Australaves and Afroaves. Although ASTRAL grouped Accipitriformes and Strigiformes as the sister to the remaining Afroaves, STEQ and wQFM-TREE supported Accipitriformes alone as sister to the remaining Afroaves consistent with concatenation analyses (5).

*Conflicting placements of Rheas* Outside of Neoaves, support for different relationships of Rheiformes within Palaeognathae was found in (5). Whereas CoalHMM (5) put Rheiformes as sister to Apterygiformes + Casuariiformes, ASTRAL-III, STEQ, and wQFM-TREE all placed Rheiformes as the sister to Tinamiformes.

### 5 Supplementary Figures and Tables

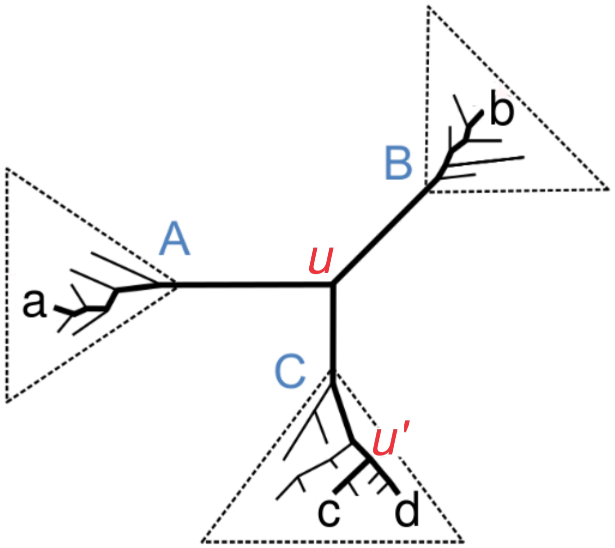


Fig. S1: Mapping quartet trees to tripartitions. Each internal node $u$ or $u'$ in the unrooted tree defines a tripartition $(A \mid B \mid C)$ of taxa. A quartet tree on $a, b, c, d$ maps to two such nodes: i) $u$, where paths from $a$ and $b$ to $c$ or $d$ converge, and ii) $u'$, where paths from $c$ and $d$ to $a$ or $b$ converge.

Table S1: Parameter settings of simulated datasets.

| Dataset | Model Parameter | Parameter Ranges |
|---|---|---|
| 37-Taxon | ILS | {0.5X, 1X, 2X } |
| | Sequence Length | {250, 500, 1000, 1500, True} |
| | Number of genes | {25, 50, 100, 200, 400, 800} |
| 48-Taxon | ILS | {0.5X, 1X, 2X} |
| | Sequence Length | {500} |
| | Number of genes | {50, 100, 200, 500, 1000} |
| 200-Taxon | Tree Length | (500K, 2M, 10M |
| | Speciation Rate | {1E-6, 1E-7} |
| 500-, 1000-Taxon | Tree Length | 2M |
| | Speciation Rate | {1E-6} |

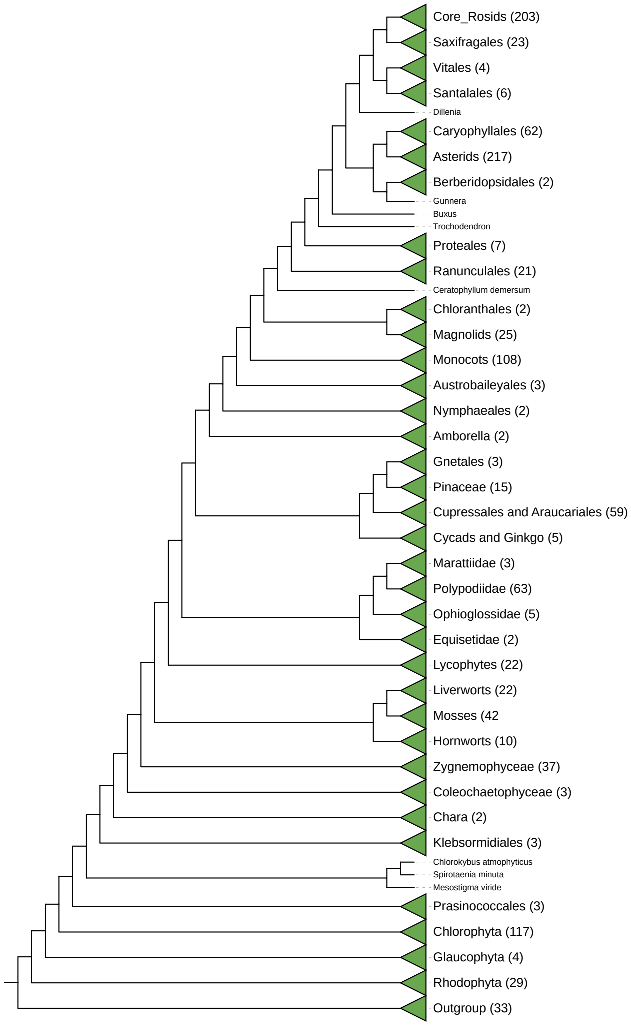


Fig. S2: Species trees estimated by wQFM-TREE using plant transcriptome dataset comprising 1,178 species and 410 gene trees

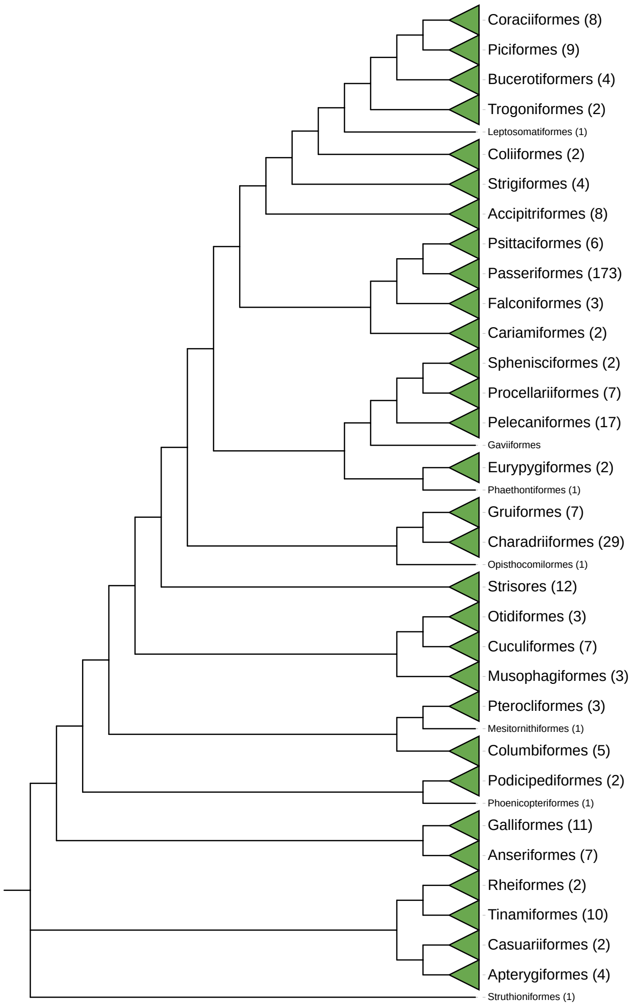


Fig. S3: Species trees estimated by wQFM-TREE using extended avian dataset comprising 363 species and 63430 gene trees (intergenic loci)

Table S2: Runtime (in seconds) of ASTRAL-III, wQFM-TREE, and STEQ on the simulated and empirical datasets. Values are shown as average (over 20 replicates for 37- and 48-taxon datasets, 10 replicates for 200- and 500-taxon datasets and 5 replicates for the 1000-taxon dataset)

| No of taxa | No of genes | ASTRAL-III | wQFM-TREE (resolved) | STEQ (resolved) |
|---|---|---|---|---|
| 37 | 200 | 1.04 | 1.2 | 0.1 |
| | 400 | 1.8 | 2.4 | 0.3 |
| | 800 | 3.9 | 5.5 | 0.5 |
| 48 | 200 | 4.8 | 3.2 | 0.3 |
| | 500 | 19.9 | 10.5 | 0.8 |
| | 1000 | 75.01 | 22.9 | 1.5 |
| 200 | 50 | 4.3 | 15.1 | 1.5 |
| | 200 | 19.8 | 76.6 | 6.1 |
| | 1000 | 297.6 | 391.1 | 30.7 |
| 500 | 50 | 24.4 | 155.3 | 10.6 |
| | 200 | 109.1 | 584.5 | 44.8 |
| | 1000 | 1509.4 | 2451.9 | 238.1 |
| 1000 | 50 | 157.5 | 898.4 | 47.6 |
| | 200 | 658.9 | 2786.8 | 200.7 |
| | 1000 | 7336.9 | 11680.4 | 1023.8 |
| Green plant data | | | | |
| 1178 | 410 | 3340.8 | 11318.7 | 422.6 |
| Extended avian data | | | | |
| 363 | 63430 | 117527.6 | 228579.5 | 12632.1 |
